# Flower visitor groups show differential responses to individual and plot-level chemodiversity with consequences for plant fitness

**DOI:** 10.1101/2023.11.21.568056

**Authors:** Rohit Sasidharan, Sean G. Grond, Stephanie Champion, Elisabeth J. Eilers, Caroline Müller

## Abstract

1. Chemodiversity, the diversity of specialised metabolites, plays a crucial role in mediating interactions between plants and animals, including insect herbivores and mutualists. Chemodiversity can be observed at both the individual and the population level. However, the impacts of chemodiversity at these two levels on interactions between plants and floral visitors, mainly pollinators and floral herbivores (florivores), are only poorly understood.
2. This study aimed to investigate the effects of chemodiversity at both individual and plot level on pollinators and florivores, examining their mutual interactions. To investigate these questions, we used individuals of the perennial *Tanacetum vulgare* differing in their terpenoid composition, representing so-called chemotypes. We planted individuals of five distinct chemotypes in a common garden design with homogeneous (five plants of the same chemotype) and heterogeneous (five different chemotypes) plots and observed flower visits in dependence of individual chemotype and plot type. Seeds were collected from a subset of plants and germination rates determined.
3. Our findings revealed that chemodiversity at the plot level significantly influenced pollinator visitation, with more visits on plants of heterogeneous plots. We also observed marginally more pollinators on one of the chemotypes grown in homogeneous plots. In contrast, chemotype but not plot type had a significant effect on florivore visits. Pollinator and florivore species richness did not vary with plot type. Furthermore, a negative correlation was observed between pollinator and florivore visits in one year, suggesting competitive interactions between these two groups. Germination rates were positively correlated with pollinator visits and affected by chemotype but not by florivore visits or plot type.
4. *Synthesis.* Our study emphasizes the significance of the scale at which different chemical profiles are perceived by flower visitors, potentially influencing the reproductive fitness of plants. Exploring the ecology of these visitors and the varying selection pressures they exert on floral chemistry can help elucidating the evolutionary processes that maintain chemodiversity in natural environments.

## 1. INTRODUCTION

Plants produce a rich repertoire of specialised (secondary) metabolites, encompassing diverse chemical classes, such as alkaloids, terpenoids or phenolics (Bennett and Wallsgrove, 1994, Dixon, 2001, Yang et al., 2018), which are involved in interactions with the environment (Wink, 2010). This metabolite composition can vary drastically not only interspecifically, but also intraspecifically (Firn and Jones, 2003, Wetzel and Whitehead, 2020), sometimes leading to distinct chemotypes. According to the interaction diversity hypothesis, chemodiversity arises from interactions with a diverse community in terms of both species and their functional roles (Kessler and Kalske, 2018, Wetzel and Whitehead, 2020). Within plants, particularly flowers engage with animals of various functional groups, aiming to attract beneficial pollinators and deter floral herbivores, known as florivores (Raguso, 2016, Sasidharan et al., 2023, Schiestl, 2015). High chemodiversity may serve these different purposes. When considering groups of plants, chemodiversity becomes more complex due to the chemical profiles of different neighbours (Salazar et al., 2016, Ziaja and Müller, 2023). Neighbouring plants may provide heterospecific but also conspecific plants with improved or reduced resistance (Bustos-Segura et al., 2017, Mutz et al., 2022, Randlkofer et al., 2010, Ziaja and Müller, 2023). Such associational effects have mostly been studied in the context of plant-(leaf)-herbivore interactions, with fewer studies on flower visitors (Salomao et al., 2006, Underwood et al., 2020). At least for pollinators, heterospecific plants may act as magnets (Underwood et al., 2020), which may likewise be the case for different intraspecific chemotypes.

Flower visitors vary in responses to floral chemodiversity. For pollinators, floral scent facilitates the location of suitable nectar or pollen resources (Raguso, 2008). Promoting effective pollen transfer enhances reproductive success for outcrossing plants (Dötterl and Vereecken, 2010). While some specialised metabolites attract pollinators, others could repel them, resulting in selective pollination and specialisation in plant-pollinator relationships (Raguso, 2008). Florivores are likewise attracted or repelled by different flower chemicals and can show preferences to specific chemotypes (Knudsen and Gershenzon, 2006, Sasidharan et al., 2023, Schiestl, 2015). Some plants avoid trade-offs between the simultaneous attraction of pollinators and florivores by modulating specialised metabolite or reward production (Kessler et al., 2013, Theis et al., 2007) or by deploying herbivore-specific repellents (Raguso, 2016). Nevertheless, interspecific competition may exist between flower visitors (Vamosi et al., 2014) and specifically between pollinators and florivores (Krupnick and Weis, 1999, Cardel and Koptur, 2010, Maloof, 2001), but not necessarily due to resource limitation (Krupnick and Weis, 1999). Feeding preferences may also shape visitor responses to chemotypes. Polyphagous insects are expected to be less influenced by unique chemotypes (Massad et al., 2022) whereas specialists might depend more on certain chemotypes and show unique preferences (Glassmire et al., 2017).

Differences in chemical composition can influence plant fitness by affecting diverse functional groups of flower visitors. Pollinators generally enhance reproductive fitness through increased pollination and seed set (Klein et al., 2007), while florivores often decrease plant fitness by reducing seed set and/or decreasing the attractiveness of flowers to pollinators (Althoff et al., 2013, McCall and Irwin, 2006). Intraspecific chemodiversity may also impact herbivory with potentially different impacts of individual chemotype- and plot-level chemodiversity (Ziaja and Müller, 2023), causing distinct plant tissue loss (Bustos-Segura et al., 2017). Furthermore, the synthesis of diverse specialised metabolites may entail certain costs associated with producing various enzymes and storage structures (Gershenzon, 1994). These factors, coupled with the genetic diversity of plants (Parker et al., 2010, Hahn et al., 2017), may result in different fitness outcomes.

*Tanacetum vulgare* L., common tansy (Asteraceae), is a perennial plant that exhibits remarkable chemodiversity in its leaf and floral terpenoid composition, forming distinct chemotypes (Keskitalo et al., 2001, Kleine and Müller, 2011). Various insects exhibited differential attraction to and/or growth performance on these chemotypes, including leaf-feeders (Ojeda-Prieto et al., 2023, Wolf et al., 2012, Ziaja and Müller, 2023) and florivores (Eilers et al., 2021, Sasidharan et al., 2023). Moreover, previous studies suggest that leaf and flower herbivores respond differently to individual chemotypes compared to the heterogeneity of chemotypes in the neighbourhood (Eilers et al., 2021, Ziaja and Müller, 2023). Such differences could lead to differences in plant fitness. Besides terpenoid chemotypes, the maternal origin (genotype) can also influence the chemodiversity of *T. vulgare* (Dussarrat et al., 2023). Thus, this species is an ideal system to study the impacts of chemodiversity on plant-animal interactions at different scales.

To investigate the effects of individual plant chemotype versus plot chemodiversity on flower visitation and plant fitness, we set up a common garden with 60 plots, half being homogenous, with five plants belonging to one of five chemotypes, the other being heterogenous, with five plants of five chemotypes. These chemotypes were defined by their leaf terpenoid profiles. Thus, we sampled leaves and flower heads of a subset of plants to determine the correlation of the terpenoid profiles between these organs. Furthermore, we scored the flower visiting insects over two years and determined the germination rate of a subset of the plants. We focused our study on pollinators and florivores, being the primary visitor groups with direct fitness effects (Junker and Blüthgen, 2010). We had the following hypotheses: (1) both pollinators and florivores show different preferences (i.e. visitation rates) on distinct chemotypes, (2) heterogeneous plots have more pollinator and less florivore visits than homogeneous plots (3) assuming interaction diversity, heterogeneous plots show a higher pollinator and florivore species diversity than homogenous plots (4) assuming competition, pollinator visits correlate negatively with florivore visits and (5) plant reproductive fitness is higher in heterogeneous plots due to higher pollinator visits.

## 2. METHODS

### 2.1 Set-up of field site

Seeds of tansy plants, growing at least 20 m apart, were collected in January 2019 from four distinct locations in Bielefeld, Germany. The seeds of these mother plants were germinated, and the terpenoid profiles of young leaves were analysed by gas chromatography-mass spectrometry (GC-MS) as outlined below (2.2). Plants of five distinct chemotypes were chosen for further investigations (Table S1 for diversity indices). Among these chemotypes, two were dominated (>55% of total terpenoid concentration) by one monoterpenoid, artemisia ketone (henceforth “Keto” chemotype) or β-thujone (“BThu”), while the other three chemotypes had two to three dominant compounds (10–50% of total terpenoid concentration), namely α-thujone and β-thujone (“ABThu”), artemisyl acetate, artemisia ketone, and artemisia alcohol (“Aacet”), or (Z)-myroxide, santolina triene, and artemisyl acetate (“Myrox”). The plants were selected from offspring of a total of 14 mother plants, with 50 plants per chemotype. From each of these stock plants, two clones were generated and planted in May 2020 in a field common garden (24 x 17 m) near Bielefeld University, Germany. Several honeybee hives stood two metres away on one side of the field. Plants were grouped into homogeneous plots containing five plants of the same chemotype (*n* = 30 plots; 6 plots per chemotype) and heterogeneous plots (*n* = 30 plots) comprising one plant from each of the five chemotypes. All plants within a single plot had distinct maternal origins. The two clones from each plant were distributed in adjacent plots of different type (i.e., homogeneous vs. heterogeneous; for specific layout details, refer to Supplementary Figure S1). Plants were placed in tubes (16 cm in diameter, 30 cm in height) inserted 25 cm deep into the soil, enabling individual plant identification (referred to as “pots” hereafter). Within plots, pots were arranged in a circular pattern with equal spacing between neighbouring plants. In 2021, a re-analysis of leaf terpenoids revealed that one plant of a homogeneous plot had been wrongly assigned. Thus, flower visitation and germination data from this plot were not included in further analyses. The plant was replaced in spring 2022 by a respective clone of the same plant to reconstitute the homogenous plot.

### 2.2 Sampling of field plants and determination of terpenoid composition

For comparison of leaf and flower head terpenoids profiles, 49 plants (8-12 per chemotype) were randomly selected from the field site. From each of these plants, the second-youngest leaf of one of the stems and two to three freshly-opened flower heads were cut, immediately frozen in liquid nitrogen and freeze-dried. Ten mg dry weight per sample was extracted with 1 mL of *n*-heptane (Roth, 99% HPLC grade), containing 10 ng µL^-1^ of 1-bromodecane (97%, Sigma Aldrich, Karlsruhe, Germany) as internal standard in an ultrasonic bath at 20 °C for 5 min (as in Eilers et al., 2021). After centrifugation, supernatants were subjected to analysis by GC-MS (GC 2010plus – MS QP2020, Shimadzu, Japan, with VF-5 MS column, 30 m x 0.2 mm inner diameter, with 10 m guard column, Varian, United States), utilising electron impact ionisation mode at 70 eV and helium as carrier gas with a flow rate of 1.5 mL min^-1^. The initial temperature of 50 °C was held for 5 min, increased to 250 °C at 10 °C per min, further to 280 °C at 30 °C per min, and held for 3 min.

Solvent blanks as controls and an alkane standard mix (C7–C40, Sigma Aldrich, Germany) were analysed using the same protocol. Terpenoids were identified by comparing the retention indices, calculated based on the alkane mix, and mass spectra with those of available synthetic reference compounds or reference data from the National Institute of Standards and Technology (NIST) 2014, Pherobase (El-Sayed, 2014) or Adams (2007). For quantification, calculated peak areas based on total ion chromatograms were divided by the sample dry weight and the peak area of the internal standard. Relative percentages were calculated based on the peak areas of all terpenoids in the sample.

### 2.3 Flower visitor observations

Flower visitor observations were performed in the flowering season (July-September) of both 2021 and 2022. Insect visitors of different species were counted at regular intervals up to 7 weeks, once per week, observing each plot for 1 min ± 10 s (about 12-20 s per plant) once in the morning and once in the afternoon of the same day (except for one afternoon count at the season end of 2021 due to rain). Flower visitors were identified to species level where possible and otherwise at least identified to genus or family level. For subsequent analyses the flower visitors were assigned functional roles, including pollinator, florivore, herbivore and predator. Arthropods sitting on the stem or below flowerheads were not counted. For further analyses, only pollinator and florivore data were considered.

The number of freshly-opened (pollen-producing, yellow) flower heads and senescing (seed-producing, brown) seed heads was estimated for each plant 1-2 d prior or after the flower visitor observations. Once during each monitoring, climate data were collected, including temperature, relative humidity (TFA Dostmann, Wertheim, Germany) and wind speed (anemometer, Voltcraft, Hirschau, Germany).

### 2.4 Determination of germination rate

Senescent flower heads were bagged using organza gauze bags (Saketos, Zittau, Germany) at the end of the flowering season of 2021 and ripe seed heads collected at the end of the year. After drying at room temperature, the seeds were separated from remaining floral structures and stored in desiccators. Germination assays were carried out only on plants of two blocks (*n* = 95 plants, excluding the plot removed from the analyses) for logistic reasons. Three replicates of approximately 35 or 100 µl of seeds per plant were spread on filter paper and pictures taken for later counting of seed numbers. Seeds were placed on glass beads (Hartenstein, diameter = 1 mm) moistened with 2 mL of deionised water in Petri dishes and stored in a climate chamber (light:dark = 16:8, 70% humidity, 20 °C). Germinated seedlings were counted and removed from the dish every 3 d until no further seeds germinated.

### 2.5 Statistical analyses

All analyses were performed using R v4.0.3 (https://www.r-project.org) and R Studio v1.4.1106 (https://posit.co/). The package *ggplot2* was used to visualise the data (Wickham, 2016).

Principal Component Analyses (PCA) were performed using the *prcomp* function from the *stats* package (R Core Team, 2018) to visualise the clustering of terpenoid profiles of leaves versus flower heads into different chemotypes. For the PCA, data were transformed using cube-root transformation, scaled using Pareto scaling and normalised to the sample medians. Correlations between the terpenoid profiles of leaves and flower heads were tested using a Mantel test with Spearman’s rank correlation method and 9999 permutations. Variation partitioning with permutational analysis of variance (ANOVA) was done with the *varpart* function in *vegan* (Oksanen et al., 2013) to explain the distribution of terpenoid variation between the chemotype and organ (i.e., flowerhead or leaf).

All flower visitor and germination data were analysed using (generalized) linear mixed effects [(G)LME] models using the packages *lme4* (Bates et al., 2015) and *glmmTMB* (for zero inflated data) (Brooks et al., 2017). Likelihood-ratio (Type III Wald chi-square) tests were used to estimate the significance of fixed effects in the package *car* (Fox, 2019) and pairwise contrast tests between levels of the fixed effects were performed with the package *emmeans* (Searle et al., 1980). Data were not transformed before implementing into the model. All models were checked using the residual diagnostic plots with the *DHARMa* package (Hartig, 2016) and models were selected based on the Akaike Information Criterion (AIC) values.

For analysis of flower visitor observations, pollinator or florivore visits were used as response variables. Chemotype and plot type were included as explanatory variables and day of sampling, wind speed, year of sampling, temperature, humidity and number of freshly-opened (pollinators) or all available flower or seed heads (florivores) were used as covariates. The block, plot, maternal origin and clonal plant identity were used as random factors. Plot was nested in block and plant identity in maternal origin. For testing correlations between pollinator and florivore visits, florivore visits were used as an explanatory variable with pollinator visits as the response variable. Correlations were analysed separately for 2021 and 2022 due to poor model convergence when combining the data. A zero-inflation term was included in the conditional models with pollinator visits as responses to account for overdispersion. Poisson or negative binomial distributions (and corresponding link functions) were assumed for these models (for details see Table 1).

**Table 1.** GLMM-derived estimates of fixed effects on (A) pollinator and (B) florivore visits. Model estimates are back-transformed from the zero-inflation corrected negative binomial distribution model. Random effects are printed in italics and given as percentages of the total variation from all effects. Significant effects (*p* < 0.05) are printed in bold, marginally significant (*p* < 0.1) in bold and italics. NT: not tested; --: not relevant; n.s.: not significant (*p* > 0.1), *: significant effect (*p* < 0.05). Keto: artemisia ketone; BThu: β-thujone; ABThu: α-thujone and β-thujone; AAcet: artemisyl acetate, artemisia ketone and artemisia alcohol; Myrox: (*Z*)-myroxide, santolina triene and artemisyl acetate.

|  | Intercept | Chemotype | Plot type | Chemotype x Plot type | Day | Wind speed | Year | Flower head number<sup>a</sup> | Temperature | Humidity | Block | Plot | Maternal origin | Plant |
| --- | --- | --- | --- | --- | --- | --- | --- | --- | --- | --- | --- | --- | --- | --- |
| **Pollinator visits: Zi-GLMM (negative binomial distribution with linear parametrization + log-link)** |  |  |  |  |  |  |  |  |  |  |  |  |  |  |
| Type III Wald Chi-square tests | $\chi^2 = 206.69$<br>df= 1<br>**$p < 0.001$** | $\chi^2 = 1.33$<br>df= 4<br>$p = 0.856$ | $\chi^2 = 5.39$<br>df= 1<br>**$p = 0.020$** | $\chi^2 = 9.17$<br>df= 4<br>**$p = 0.057$** | $\chi^2 = 1654.84$<br>df= 1<br>**$p < 0.001$** | $\chi^2 = 2.29$<br>df= 1<br>$p = 0.130$ | $\chi^2 = 206.87$<br>df= 1<br>**$p < 0.001$** | $\chi^2 = 290.59$<br>df= 1<br>**$p < 0.001$** | $\chi^2 = 3.87$<br>df= 1<br>**$p = 0.049$** | $\chi^2 = 12.39$<br>df= 1<br>**$p < 0.001$** | -- | -- | -- | -- |
| Conditional model fixed effects levels or random effects variances | -- | AAcet – Intercept = - 0.008, n.s.<br>BThu – Intercept = 0.09, n.s.<br>Keto – Intercept = 0.08, n.s.<br>Myrox – Intercept = 0.01, n.s. | Homogeneous – Intercept = **- 0.21 \*** | AAcet homogeneous – Intercept = **0.03 \***<br>BThu homogeneous – Intercept = 0.04 n.s.<br>Keto homogeneous – Intercept = 0.03, $p = 0.07$<br>Myrox homogeneous – Intercept = **0.03 \*** | -- | -- | -- | -- | -- | -- | 1.02 % | 0.66 % | 1.39 % | 6.17 % |
| Zero-inflation model | Estimate = - 2.42<br>**$p < 0.001$** | NT | NT | NT | Estimate = -0.26<br>**$p = 0.057$** | NT | NT | NT | NT | NT | NT | NT | NT | NT |
| **Florivore visits: Zi-GLMM (Poisson distribution + log-link)** |  |  |  |  |  |  |  |  |  |  |  |  |  |  |
| Type III Wald Chi-square tests | $\chi^2 = 20.68$<br>df= 1<br>**$p < 0.001$** | $\chi^2 = 10.55$<br>df= 4<br>**$p = 0.032$** | $\chi^2 = 0.013$<br>df= 1<br>$p = 0.911$ | $\chi^2 = 2.61$<br>df= 4<br>$p = 0.625$ | $\chi^2 = 553.04$<br>df= 1<br>**$p < 0.001$** | NT | $\chi^2 = 20.66$<br>df= 1<br>**$p < 0.001$** | $\chi^2 = 14.90$<br>df= 1<br>**$p < 0.001$** | $\chi^2 = 18.04$<br>df= 1<br>**$p < 0.001$** | $\chi^2 = 553.04$<br>df= 1<br>**$p < 0.001$** | -- | -- | -- | -- |
| Conditional model fixed effects levels | -- | AAcet – Intercept = - 0.1833 n.s.<br>BThu – Intercept = | Homogeneous – Intercept = - 0.0133 n.s. | AAcet Homogeneous – Intercept = - 0.0642 n.s. | -- | -- | -- | -- | -- | -- | 0.61 % | 0.45 % | 0.27 % | 5.77 % |
| or random effects variances |  | - 0.0575 n.s.<br>Keto – Intercept = - 0.0744 n.s.<br>Myrox – Intercept = 0.272<br>***p* = 0.073** |  | BThu Homogeneous – Intercept = - 0.0842 n.s.<br>Keto Homogeneous – Intercept = 0.1265 n.s.<br>Myrox Homogeneous – Intercept = - 0.1151 n.s. |  |  |  |  |  |  |  |  |  |  |
| Zero-inflation model | Estimate = 1.402<br>***p* < 0.001** | NT | NT | NT | NT | NT | NT | Estimate = -0.040<br>***p* < 0.001** | NT | NT | NT | NT | NT | NT |

Differences in insect species richness in dependence of plot type were tested using the number of species belonging to either pollinators or florivores present within each plot. Plot type was used as a fixed effect and block and plot were used as random factors in a linear mixed effects model. Day of sampling, wind speed, year of sampling, temperature, humidity and number of fresh flower heads were included as covariates.

Germination rate (%) as the response variable was analysed with a linear model (LM), with plot type and chemotype as fixed effects. An ANOVA followed by Tukey’s pairwise comparisons were used to analyse significant differences among and between factor levels. For testing the correlation between germination rate and pollinator or florivore visits, a linear mixed effects model was used, with germination rate as response variable and pollinator or florivore visits as well as plot type as explanatory variables. Numbers of fresh yellow flower heads only (pollinator) or all available flower and seed heads per plant (for florivores) were used as covariates. The chemotype, block, plot, maternal origin and plant identity were included as random factors.

## 3 RESULTS

### 3.1 Terpenoid profiles of leaves versus flower heads

Overall, 77 terpenoids were detected in leaves and 79 in flower heads. The relative terpenoid composition of the leaves was mostly mirrored in the flower heads, with the dominant terpenoids determining the profiles of both (Figure 1A). The five chemotypes could be discriminated based on their terpenoid profiles in both leaves and flower heads (Figure 1 B,C), indicating that terpenoid profiles are chemotype-specific in both organs. A Mantel test revealed a significant correlation between the two profiles (Mantel’s *r* = 0.82, Spearman’s rank correlation, *p* < 0.001, *n_permutation_*= 9999). Chemotype explained a large part of the terpenoid variation among samples when compared to the organ (Figure 1D).

**Figure 1.**
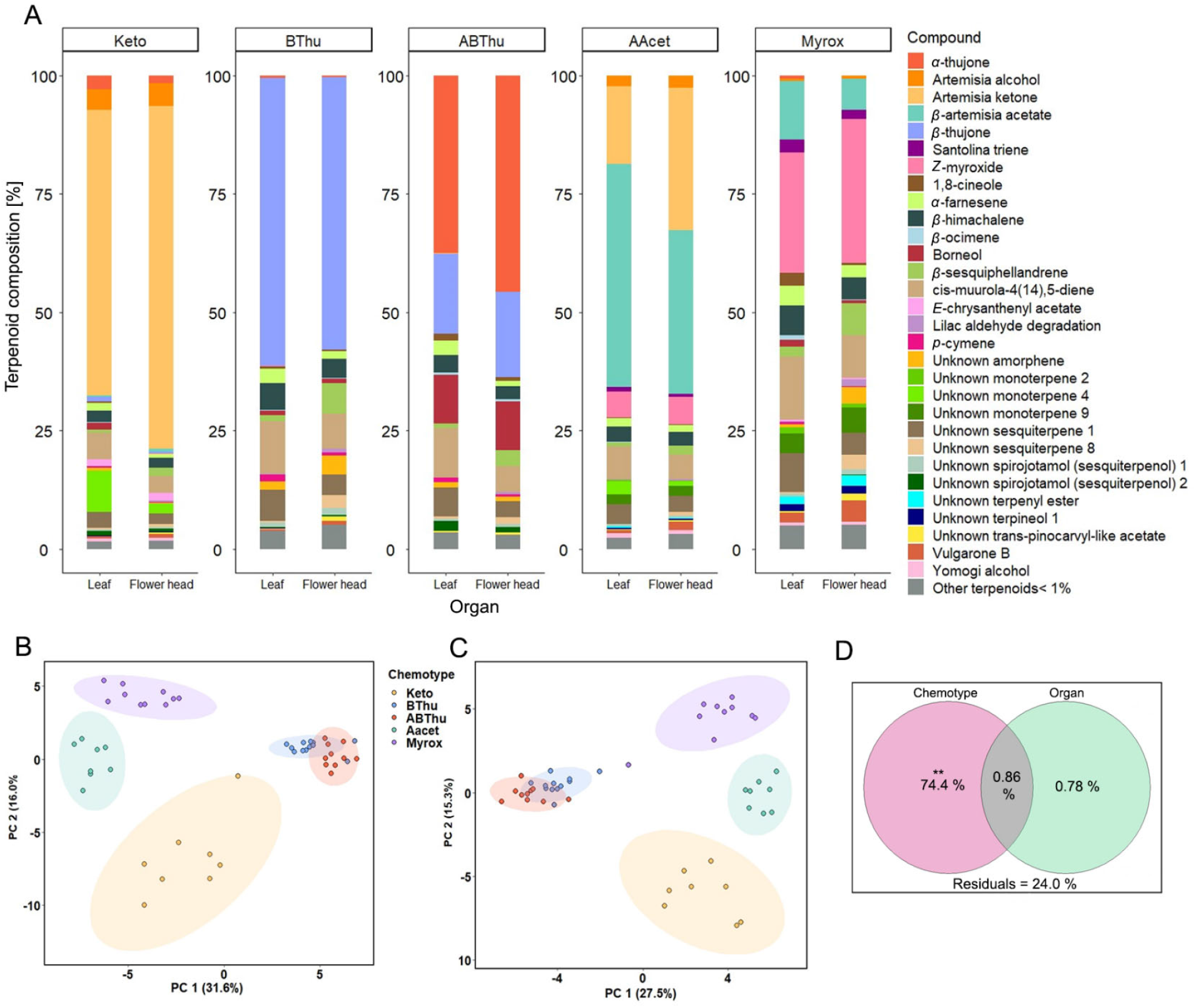
Terpenoid composition of leaves and flower heads of *Tanacetum vulgare* plants (*n* = 49 individuals, 8-12 per chemotype) sampled in the experimental field in 2022. (A) Mean relative composition (A), PCA scores plot of (B) leaf and (C) flower head terpenoids and (D) variation partitioning of terpenoid profile between chemotype and organ (i.e., leaf or flower head) (permutational ANOVA on marginal models, *n_permutation_* = 999, *F*_chemotype_ = 62.003, *F*_organ_ = 1.348, **: *p* < 0.01). Chemotypes were defined by their main terpenoids; Keto: artemisia ketone; BThu: β-thujone; ABThu: α-thujone and β-thujone; Aacet: artemisyl acetate, artemisia ketone and artemisia alcohol; Myrox: (*Z*)-myroxide, santolina triene and artemisyl acetate.

### 3.2 Differences in flower visitations by pollinators and florivores

Pollinator, florivore and overall insect counts over the flowering season were much higher in 2021 compared to 2022 (Table S2). In both years together, the number of observed pollinators was roughly five times higher than the number of florivores. As potential pollinators, mostly hymenopterans including *Apis mellifera*, wild bees (*Andrena* spp.) and dipterans were observed, with *A. mellifera* far outnumbering the other species. The observed pollinators are believed to mainly be generalists, visiting different plant species (Table S2). Pollinator visits were not influenced by chemotype alone, but differed depending on the plot type and marginally on the interaction of chemotype and plot type (Table 1). On plants growing in heterogeneous plots, more pollinators were estimated than in homogenous plots (*χ²* = 5.39, *p* = 0.02). However, plants of the Aacet and Myrox chemotypes had more pollinators when growing in homogeneous than in heterogenous plots (Table 1, Figure S2). Responses of *A. mellifera* were similar to those of overall pollinators, but the interactive effect of plot type and chemotype was significant for this species (Table S3).

The main florivores were coleopterans, namely phalacrids (*Olibrus* spp.) and cantharids, and some hemipterans. Beetles of the genus *Olibrus* spp. were the most dominant florivores and are known specialists on *T. vulgare* or related Asteraceae (Gaafar et al., 2016, Müller-Schärer and Brown, 1995) (Table S2). Florivore visits differed significantly between chemotypes (Table 1), with on average most florivores found on plants of the Myrox chemotype. Contrary to the pollinator visits, the florivore visits were not affected by plot type or the interaction of chemotype and plot type (Table 1, Figure S2). In 2021, there was a negative correlation between pollinator and florivore visits (*χ²* = 3.91, *p* = 0.048; Figure 2A). However, this effect was not observed in 2022 (*χ²* = 0.17, *p* = 0.676; Figure S2). *Olibrus* spp. were not influenced by chemotype or plot type in the field, unlike overall florivores (Table S3).

**Figure 2.**
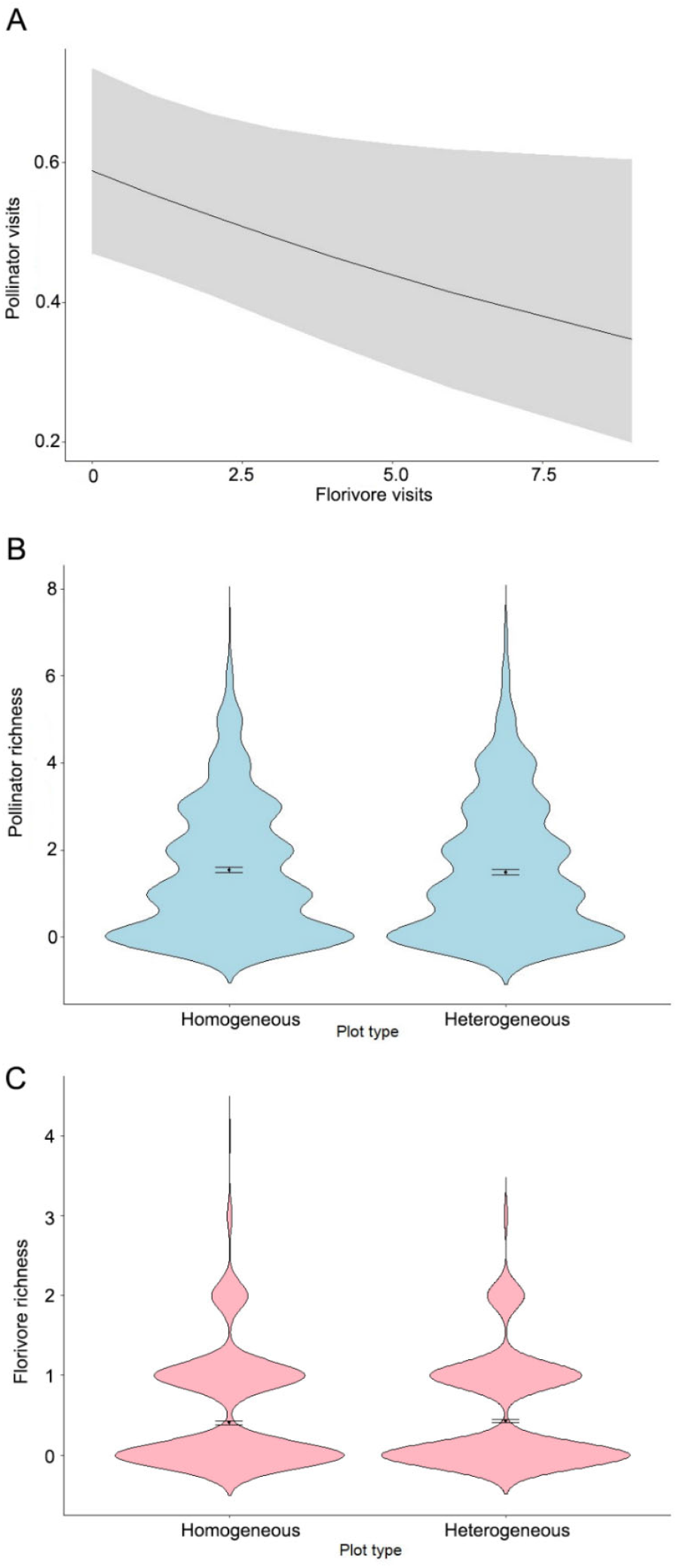
(A) Correlations (model-estimated) between pollinator and florivore visits in 2021 based on a zero-inflation negative-binomial distribution model. Violin plots showing the plot-level species richness of (B) pollinators and (C) florivores on homogenous and heterogeneous plots of different chemotypes of *Tanacetum vulgare*. No significant differences were found between plot types. Model-derived means ± SEs are also shown (LME).

Random effects, including block, plot and maternal origin did not explain much of the variation in either pollinator or florivore visits. Plant identity, however, accounted for a high amount of variation (>5%) in the visits.

The species richness of pollinators (Type III Wald chi-square tests, *χ²* = 0.97, *p* = 0.324) and florivores did not differ between plot types (*χ²* = 0.67, *p* = 0.415).

### 3.3 Germination rates of field-collected seeds in relation to flower visitors

The germination rates of plants collected from two of the six plots were significantly affected by chemotype (LM, *F* = 3.45, *p* = 0.009) but not by plot type (*F* = 0.001, *p* = 0.974). They were significantly lower on plants of the ABThu and the Aacet chemotype and marginally lower on plants of the BThu chemotype compared to the Myrox chemotype (Tukey’s pairwise comparisons) (Figure 3A). The germination rates were positively correlated with the pollinator visits (LME, *F* = 4.75, *p* = 0.030; Fig 4A) but not with florivore visits (*F* = 0.81, *p* = 0.370; Figure 4B).

**Figure 3.**
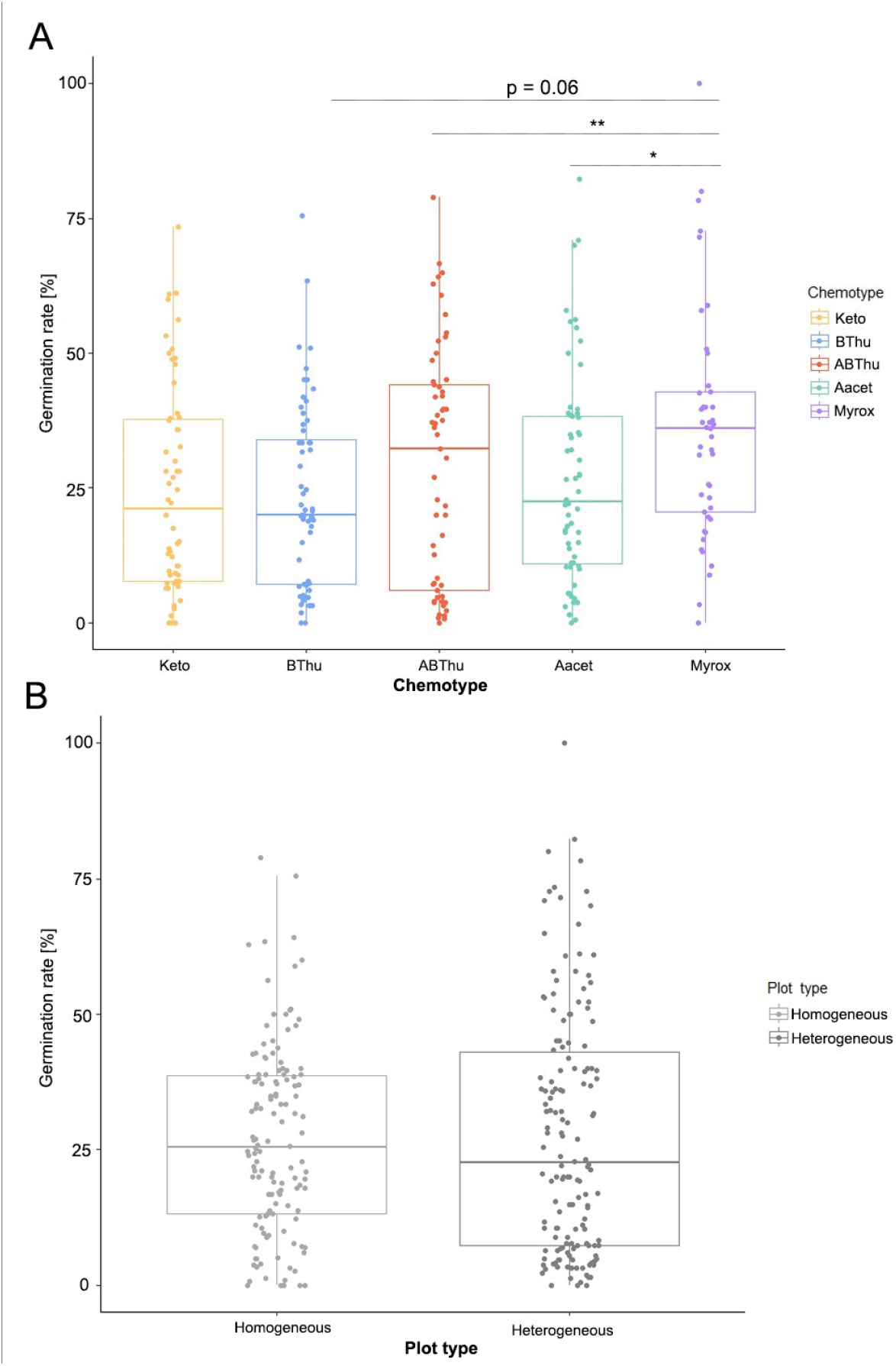
Germination rates of seeds of field-grown *Tanacetum vulgare* plants of (A) different chemotypes (Keto: artemisia ketone; BThu: β-thujone; ABThu: α-thujone and β-thujone; Aacet: artemisyl acetate, artemisia ketone and artemisia alcohol; Myrox: (*Z*)-myroxide, santolina triene and artemisyl acetate) and (B) different plot types. Box plots show median, lower (Q1) and upper (Q3) quartiles with whiskers, i.e., 1.5 times the inter-quartile range (Q3-Q1). Asterisks indicate a significant difference (*p* < 0.05) in pairwise contrast tests between model-derived means. Three replicates were used for each plant individual (*n* = 95 plants) and replicates were included as a covariate in the mixed model.

**Figure 4.**
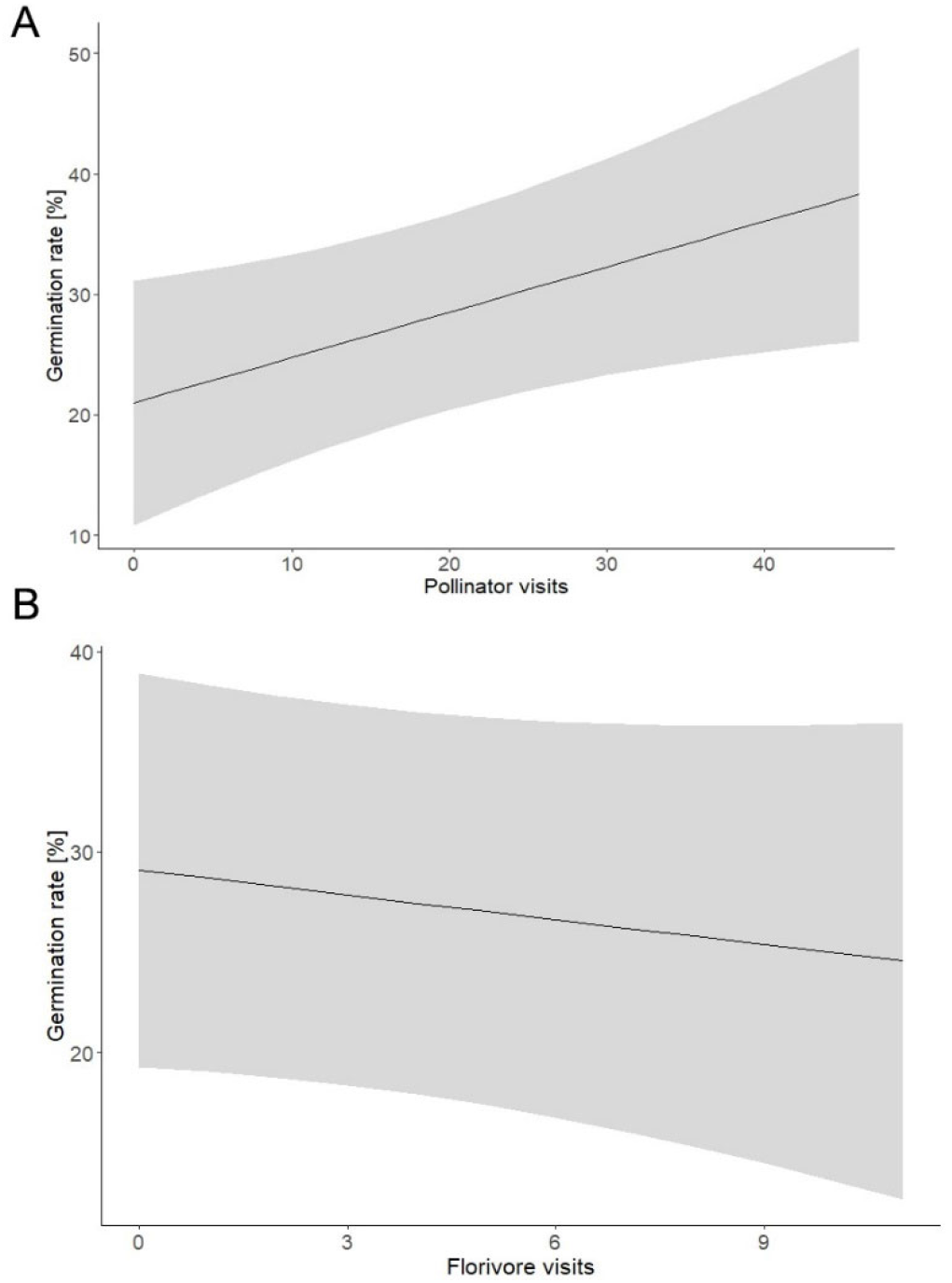
Correlation of germination rates of *Tanacetum vulgare* seeds with either (A) pollinator or (B) florivore visits. Model estimates are back-transformed from the linear mixed effects model. Three replicates were used for each clonal plant and were included as a covariate in the mixed model (*n* = 95 plants). The relation was significant for pollinator visits (LME, *F* = 4.75, *p* = 0.030) but not for florivore visits (*F* = 0.81, *p* = 0.370).

## DISCUSSION

The chemodiversity of individual plants and their associated neighbourhoods are expected to have an influence on insect visitations (Kessler and Kalske, 2018, Salazar et al., 2016, Underwood et al., 2020, Ziaja and Müller, 2023), yet this is not well-studied for floral mutualistic and antagonistic visitors. The results from our field study on the chemodiverse plant species *T. vulgare* show that individual (i.e. chemotype) as well as plot-level diversity (i.e. plot type) had differential effects on pollinators and florivores, with overall more pollinators visiting chemically heterogeneous plots, whereas florivore distribution depended on the individual plant chemotype within these plots (Figure 5). Since pollinators correlated positively with the germination rate, they may have affected plant fitness, although the germination rate also depended on the chemotype. Species diversity of pollinators and florivores at the plot level was not affected by the plot type.

**Figure 5.**
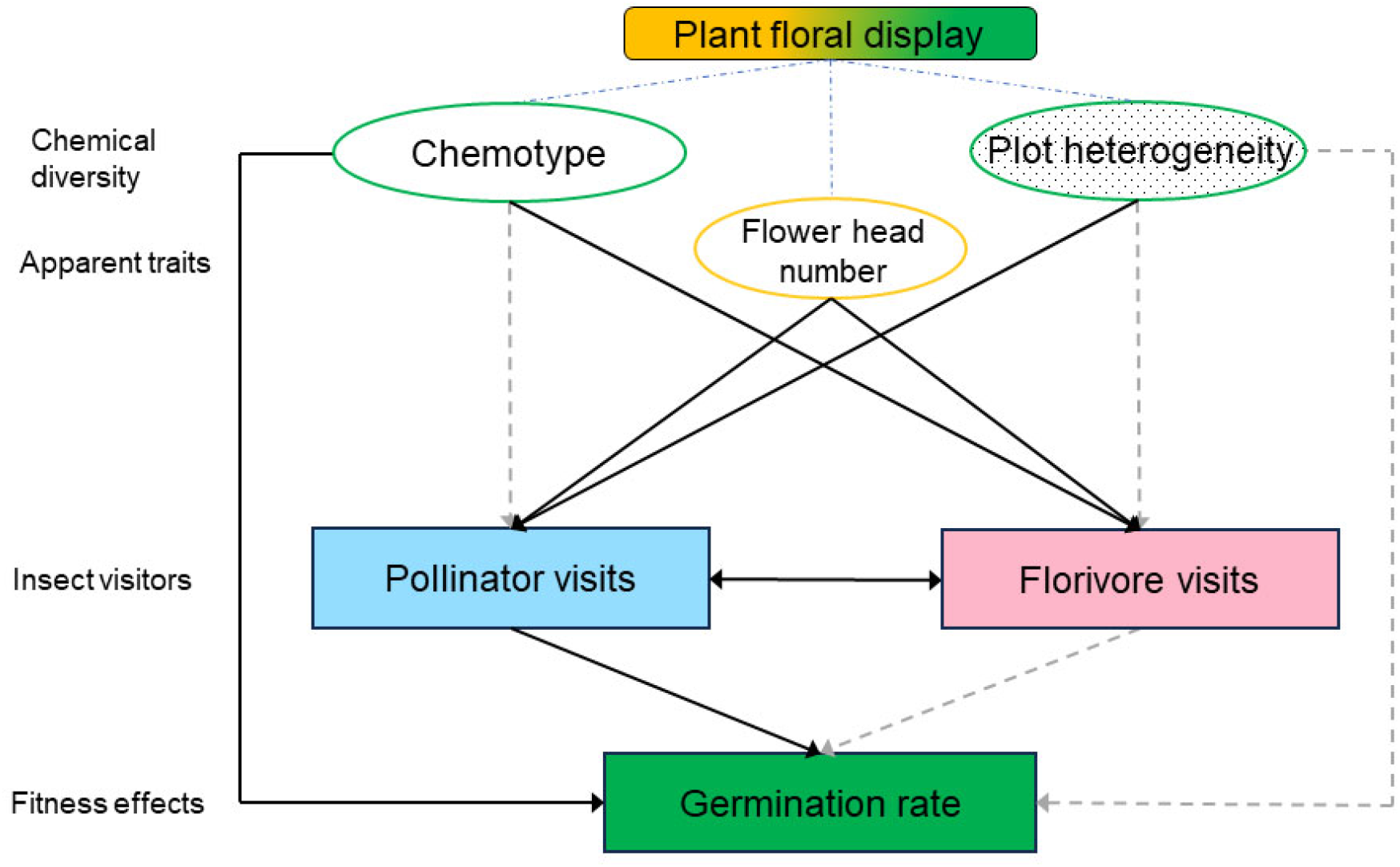
Schematic representation of the role of chemotype and plot type in the field and their influences on flower visitors. These flower visitors may in turn influence plant reproductive fitness directly or indirectly. Black continuous lines indicate significant relations, while dashed grey lines indicate non-significant relations.

The flower heads showed the same chemotype as the leaves in our field-grown plants, as previously also found for *T. vulgare* plants grown under a controlled greenhouse environment (Sasidharan et al., 2023). Thus, in this species the chemotype, defined here by one to three dominant terpenoids, is a unique phenotype of the individuals’ aboveground organs and consistent under different conditions (Kleine and Müller, 2011, Benedek et al., 2015, Ziaja and Müller, 2023). A correlated pattern in leaves and flower heads may minimise the costs that exist for terpenoid biosynthesis (Gershenzon, 1994). Nevertheless, overall terpenoid concentrations and composition can plastically respond to environmental challenges in *T. vulgare* (Clancy et al., 2016, Kleine and Müller, 2013, Kleine and Müller, 2014).

The chemotype impacted pollinators and florivores in distinct ways. For pollinators, the effect of chemotype was only marginally significant in interaction with plot type. In contrast, florivore visits differed between chemotypes, in line with our hypothesis (1), with most florivores found on the Myrox chemotype. This may imply that herbivores, including florivores, are one of the primary targets of aboveground chemotype-defining terpenoids. The major florivore observed in our study was *Olibrus* spp. (Table 1), which are specialists on Asteraceae (Gaafar et al., 2016). Preferences for certain chemotypes were also found in laboratory studies for *Olibrus aeneus*, which preferred the BThu over the Myrox chemotype in olfactometer and contact dual choice assays (Sasidharan et al., 2023). In the field, preferences may be related to different abundances of certain chemotypes (Bálint et al., 2016, Bustos-Segura et al., 2015, Humay et al., 2023) and depend on local adaptations of insect populations (Laukkanen et al., 2012, Kalske et al., 2016). The florivores observed in our study comprised several genera (Table 1), which may explain distinct preferences, but the main florivores in our field study, *Olibrus* spp., were not influenced by chemotype when tested separately, possibly due to confounding effects such as competition or low numbers (Table S3). Different florivore species have been shown to distinguish between chemically-different flowers of cultivated and wild species (Egan et al., 2018, Eilers et al., 2021, Kessler et al., 2013, Rusman et al., 2022).

Pollinators are believed to be dietary generalists when compared to herbivores (Fontaine et al., 2009, Jones and Agrawal, 2017). Polyphagous insects could be less affected by chemotypes compared to dietary specialists. Specialist herbivory decreased but generalist herbivory was not impacted by chemical diversity in *Piper* spp. (Massad et al., 2022). Specialists are often more susceptible to chemotypic or genotypic variation, which could affect their performance or community structure (Glassmire et al., 2017, Hervé et al., 2016). Florivores are also likely not as mobile as generalist pollinators (Jones and Agrawal, 2017, Underwood et al., 2020) and may be more selective to certain chemotypes. The correct choice of a suitable chemotype should thus be critical to their survival and that of their progeny (Gripenberg et al., 2010).

In line with our hypothesis (2), we observed more pollinators on heterogeneous than on homogeneous plots. Unlike florivores, pollinators may be particularly attracted to chemically diverse patches. Pollinators such as bees often mix dietary sources to optimise their nutrition (Bukovinszky et al., 2017). At the same time, dietary mixing can lead to a dilution of individual “toxins” (Eckhardt et al., 2014, Friedrichs et. al, 2022). In small doses, several toxins may physiologically protect bee colonies against pathogens and diseases (Spear et al., 2016, Manson et al., 2010). Moreover, as flying, rapidly-foraging insects, pollinators may perceive the floral blend of a patch of plants in direct neighbourhood, while the less mobile florivores may rather perceive individual plant headspace odours. Thus, exhibiting multiple chemotypes, as observed in natural settings for *T. vulgare* (Clancy et al., 2016, Kleine and Müller, 2011) can be advantageous towards attracting more pollinators, leading to higher pollination and fitness. We did not observe a trade-off in attracting more florivores in heterogeneous plots nor did we find evidence for associational resistance of these plots towards florivores. Contradictory to our hypothesis (3), pollinator and florivore diversity was not higher in heterogenous than in homogenous plots. Plot-level chemodiversity did not show an influence on interaction diversity (Kessler and Kalske, 2018, Wetzel and Whitehead, 2020). Plot type-driven increase in pollinator counts was therefore simply due to more pollinators rather than more interactions with a diverse pollinator community.

In line with our hypothesis (4), pollinator and florivore visits in this study showed a negative correlation. This finding suggests competitive interactions between these visitor groups. A negative impact of florivory on pollinators without direct damage of pollen or seeds has been reported in other systems (Cardel and Koptur, 2010, McCall and Irwin, 2006, Althoff et al., 2013). However, we observed many more pollinators than florivores, likely due to *A. mellifera* hives present near the field site. Competition between the two groups might also suggest that under our field conditions, pollinators minimised the influence of florivores, which may not be the case for other settings. Due to their low numbers, we did not consider the effects of predators and leaf herbivores on species interactions.

Chemodiversity is suggested to have an effect on plant fitness (Bustos-Segura et al., 2017, Salazar et al., 2016, Wetzel and Whitehead, 2020), for example due to associational resistance (Bustos-Segura et al., 2017, Hauri et al., 2022). We scored the germination of a subset of seeds collected from our field plants as a fitness proxy. Although we had expected heterogeneous plots to show a higher fitness (hypothesis 5), we did not find such an effect. Only individual plant chemotype influenced germination. Seeds from the Myrox chemotype, despite having more florivores, also had more pollinators when present in homogeneous plots, and showed a higher germination rate than some of the other chemotypes (Figure 3). Because *T. vulgare* is self-incompatible (Lokki et al., 1973), outcrossing mediated by pollinators is essential for a viable seed set. Intuitively, over all studied plants, germination was positively correlated with pollinator visits. Positive relationships between germination rate and pollinator visits have been found in many species, depending on the pollinator-to-flower ratio (Lázaro et al., 2013, Waser and Price, 1983). In contrast, florivores are known to negatively impact plant fitness (Althoff et al., 2013, Cardel and Koptur, 2010, Cascante-Marín et al., 2008). In our study there was by trend a negative relation, which, however, was not significant. Florivore visitation is thus not necessarily unfavourable for plants, especially if compensated by higher pollinator visits. Differences in germination rate among chemotypes also point to intrinsic effects, which may be related, for example, to differential biosynthetic costs of terpenoid production (Kleine and Müller, 2013, Neilson et al., 2013). Moreover, some terpenoids can act as antimicrobial defences (Tiku, 2018), potentially helping in the germination process, which is sensitive to attack (Crist and Friese, 1993)

Summarising our study, chemodiversity proved to be important at the individual plant chemotype level towards florivores and at the plot level towards pollinators. Only minor influences of the maternal origin or plant identity were found for pollinator or florivore visits (random variances, Table 1), so any observed differences are likely based on chemodiversity. Spatial diversity in chemical composition may be an important driver of herbivore rather than pollinator specialisation (Becerra, 2007, Richards et al., 2015). In contrast, being dietary generalists, pollinators may broaden their use of resources (Burkle et al., 2013) without specializing on certain chemotypes. The resulting effects on reproductive fitness of the plants likely originated from different contributing factors such as pollinator visitation, florivore-related damage and biosynthetic costs of chemical diversity. Chemodiversity was found to directly influence flower visitor numbers, without leading to more interactions with a higher diversity of species, thus challenging the interaction diversity hypothesis (Kessler and Kalske, 2018). Finally, a key assumption in florivore-related studies is that the florivores usually damage the flowers or reduce pollination. This may not always be the case as shown in a few studies (Burkle et al., 2007, Tsuji et al., 2016). Instead, florivores may even transfer pollen between plants and contribute to pollination service (McCall and Irwin, 2006, Etl et al., 2022). More research is necessary on the effects of chemodiversity on different flower visitors under natural settings to fully understand the processes shaping these observations.

## AUTHOR CONTRIBUTIONS

EJE and CM developed the research questions and study design. RS, SC and EJE developed the methodology. All authors contributed towards acquiring the data. RS and SG analysed the data and RS wrote the first version of the manuscript. RS and CM led the writing of the manuscript and all authors revised it.

## ACKNOWLEDGEMENTS

We thank Dominik Ziaja, Lukas Brokate, Ruth Jakobs and Tanja Bloss for practical help, and Gebhard Sewing and members of the Chemical Ecology department for help with maintaining the field setup. We also thank Karin Schrieber, Silvia Eckert and Thomas Dussarrat for help with statistical analyses. This research was funded by the German Research Foundation (Deutsche Forschungsgemeinschaft, DFG), as part of the research unit FOR 3000 (EI 1164/1-1).

