## Supplement for "Flower visitor groups show differential responses to individual and plot-level chemodiversity with consequences for plant fitness"

**Supplementary Information**

**TABLES**

**Table S1**. Shannon diversity ($H_s$) and richness (means ± SD) of terpenoid profiles of leaves and flowers collected in the field from *Tanacetum vulgare* of five chemotypes.

| **Chemotype** | **Leaf** | | **Flower head** | |
|---|---|---|---|---|
| | $H_s$ | richness | $H_s$ | richness |
| Keto | 1.48 ± 0.24 | 35.38 ± 2.77 | 1.18 ± 0.35 | 35.25 ± 4.59 |
| BThu | 1.54 ± 0.24 | 34.5 ± 3.48 | 1.58 ± 0.32 | 32.92 ± 4.34 |
| ABThu | 1.85 ± 0.13 | 37.5 ± 2.99 | 1.65 ± 0.19 | 33.8 ± 2.70 |
| Aacet | 1.94 ± 0.18 | 42.22 ± 1.72 | 1.85 ± 0.49 | 39.78 ± 5.19 |
| Myrox | 2.48 ± 0.24 | 39.3 ± 4.60 | 2.39 ± 0.40 | 36.1 ± 5.86 |

**Table S2.** Distribution of flower visitors for both years across functional roles, orders and taxa, with dietary specialisations provided for pollinators or florivores, if known.

| Functional role | Order | Taxon | Comment | Functional role and specialisation (with reference) | | 2021 | 2022 | Grand total |
|---|---|---|---|---|---|---|---|---|
| Pollinator | Diptera | Fly | Mix, mainly *Musca* spp. | Generalist | doi:10.1177/00368504231184035, doi: 10.3390/insects11060341; pollinates several families of plants | 187 | 536 | 349 |
| Pollinator | Diptera | Fruit moths | Mix/unknown | Unknown | -- | 86 | 86 | 0 |
| Pollinator | Diptera | Green bottle fly | *Lucilia sericata* | Generalist | doi: 10.3390/insects11060341, doi: 10.1098/rspb.2002.2174 | 95 | 291 | 196 |
| Pollinator | Diptera | Mosquito | Unknown | Generalist | doi.org/10.1111/eea.12852 | 3 | 3 | 0 |
| Pollinator | Diptera | Other moths | Unknown | Unknown | doi: 10.1007/s11829-016-9414-3 | 1 | 1 | 0 |
| Pollinator | Diptera | *Myoleja* spp. | Unknown | Unknown | -- | 12 | 29 | 17 |
| Pollinator | Diptera | Syrphid | Known generalists | Generalist | doi: 10.1016/j.cub.2022.03.028, doi: 10.3390/insects11060341 | 109 | 427 | 318 |
| Pollinator | Hymenoptera | *Andrena* spp. | Known generalists | Generalist or specialist | https://www.jstor.org/stable/26453772 | 4 | 620 | 616 |
| Pollinator | Hymenoptera | *Apis mellifera* | Known generalists | Generalist | https://www.jstor.org/stable/90009239 | 4420 | 5274 | 854 |
| Pollinator | Hymenoptera | *Athalia* | Known generalists on non-Brassicaceae | Generalist | doi/10.1111/mec.16943; high network links to various plant species | 0 | 1 | 1 |
| Pollinator | Hymenoptera | Bumblebee (*Bombus* spp.) | Known generalists | Generalist, rarely specialist | https://www.xerces.org/bumblebees/about doi: 10.3389/fevo.2021.699649 | 0 | 7 | 7 |
| Pollinator | Hymenoptera | *Colletes* spp. | Known generalists | Generalist, occasionally specilaist | doi: 10.1111/nph.13016 | 354 | 354 | 0 |

| | | | | | | | | |
|---|---|---|---|---|---|---|---|---|
| Pollinator | Hymenoptera | Masked bee | Mainly *Hylaeus* spp. | Generalist | doi: 10.1371/journal.pone.0088948 | 8 | 38 | 30 |
| Pollinator | Hymenoptera | Ruby tail wasp | Mainly *Chrysis* spp. | Unknown | doi: 10.5962/bhl.title.21626; Pollinator interactions between Chrysis sp. and Rosa canina | 4 | 5 | 1 |
| Pollinator | Hymenoptera | Sweat bee | *Halictus* or *Lasioglossum* spp. | Generalist | https://nativebeeology.com/mind-your-bees-and-gardens-2/sweat-bee/ | 164 | 231 | 67 |
| Pollinator | Hymenoptera | Wasp | *Vespula* or *Dolichovespula* spp. | Generalist | doi: 10.3389/fevo.2020.571454 | 10 | 66 | 56 |
| Pollinator | Lepidoptera | Copper butterfly | *Lycaena* spp. | Varied | doi: 10.1111/icad.12087; depends on range | 7 | 8 | 1 |
| Pollinator | Lepidoptera | Other lycaenids | *Polyommatus* or *Phengaris* spp. | Unknown | https://www.pollinator.org/pollinator.org/assets/generalFiles/Lepidoptera-Fact-Sheet.pdf | 1 | 3 | 2 |
| Pollinator | Lepidoptera | Pieris | *Pieris rapae* | Generalist | doi: 10.1098/rspb.2002.2174 | 3 | 3 | 0 |
| Total pollinators | | | | | | **5468** | **2515** | **7983** |
| Florivore | Coleoptera | Cantharid (others) | Mix/Unknown | Generalist | https://tuprints.ulb.tu-darmstadt.de/id/eprint/5458 | 12 | 12 | 0 |
| Florivore | Coleoptera | Chrysomelid | *Chrysolina graminis* | Specialist | Chapman, Dan. "Spatial ecology of the tansy beetle (Chrysolina graminis)." (2006). | 0 | 1 | 1 |
| Florivore | Coleoptera | Coccinellid | Mainly *Coccinella septumpunctata* | Generalist | doi: 10.14411/eje.2005.076 | 35 | 77 | 42 |
| Florivore | Coleoptera | Mordellid | Mainly *Mordella* spp. | Generalist | https://tuprints.ulb.tu-darmstadt.de/id/eprint/5458 | 31 | 43 | 12 |
| Florivore | Coleoptera | Oedemerid | Mix/Unknown | Unknown | https://tuprints.ulb.tu-darmstadt.de/id/eprint/5458 | 40 | 42 | 2 |
| Florivore | Coleoptera | Phalacrid | Primarily *Olibrus* spp. | Specialist | In main text | 579 | 1134 | 555 |
| Florivore | Coleoptera | Seed beetle | Mix/Unknown | Unknown | Feeds on seeds and seed heads | 2 | 2 | 0 |
| Florivore | Coleoptera | Soldier beetle | Mainly *Rhagonycha fulva* | Generalist | https://tuprints.ulb.tu-darmstadt.de/id/eprint/5458 | 198 | 329 | 131 |
| Florivore | Hemiptera | Pyrrhocorid | *Pyrrhocoris* or | Malvaceae | doi: 10.1038/ismej.2015.75 | 30 | 30 | 0 |

| | | | | | | | | |
|---|---|---|---|---|---|---|---|---|
| | | | *Corizus sps.* | specialists, facultative generalists | | | | |
| Total florivores | | | | | | **927** | **743** | **1670** |
| Herbivore | Coleoptera | Jewel beetle | -- | -- | | 2 | 2 | 0 |
| Herbivore | Hemiptera | Aphid | -- | -- | | 0 | 14 | 14 |
| Herbivore | Hemiptera | Cicadid | -- | -- | | 0 | 14 | 14 |
| Herbivore | Hemiptera | Jewel shield bug | -- | -- | | 80 | 88 | 8 |
| Herbivore | Hemiptera | Leaf foot bug | -- | -- | | 0 | 15 | 15 |
| Herbivore | Hemiptera | Leaffoot bug | -- | -- | | 135 | 135 | 0 |
| Herbivore | Hemiptera | Mirid bug | -- | -- | | 468 | 816 | 348 |
| Herbivore | Hemiptera | Shield bug | -- | -- | | 60 | 121 | 61 |
| Herbivore | Hemiptera | Stick bug | -- | -- | | 3 | 3 | 0 |
| Herbivore | Lepidoptera | Lepidopteran larva | -- | -- | | 2 | 2 | 0 |
| Herbivore | Orthoptera | Tettigonia | -- | -- | | 2 | 3 | 1 |
| Herbivore | Unknown | Weevil | -- | -- | | 0 | 4 | 4 |
| Total herbivores | | | | | | **752** | **465** | **1217** |
| Predator | Araneae | Spider | -- | -- | | 98 | 187 | 89 |
| Predator | Crabronidae | Beewolf | -- | | | 0 | 5 | 5 |
| Predator | Diptera | Predatory flies | -- | -- | | 0 | 7 | 7 |
| Predator | Hemiptera | Damsel bug | -- | -- | | 5 | 22 | 17 |
| Predator | Hymenoptera | Ant | -- | -- | | 7 | 11 | 4 |
| Predator | Hymenoptera | Hornet | -- | -- | | 0 | 2 | 2 |

| | | | | | | | | |
|---|---|---|---|---|---|---|---|---|
| Predator | Hymenoptera | Parasitoid | -- | -- | | 0 | 108 | 108 |
| Predator | Neuroptera | Chrysopid | -- | -- | | 15 | 15 | 0 |
| Total predators | | | | | | **125** | **232** | **825** |
| **Sum total of all insects** | | | | | | **7272** | **3955** | **11227** |

**Table S3.** GLMM-derived estimates of fixed effects on (A) *Apis mellifera* and (B) phalacrid (*Olibrus* spp.) visits. Model estimates are back-transformed from the zero-inflation corrected negative binomial distribution model. Random effects are printed in italics and given as percentages of the total variation from all effects. Significant effects ($p < 0.05$) are printed in bold, marginally significant ($p < 0.1$) in bold and italics. NT: not tested; --: not relevant; n.s.: not significant ($p > 0.1$), *: significant effect ($p < 0.05$). Keto: artemisia ketone; BThu: β-thujone; ABThu: α-thujone and β-thujone; Aacet: artemisyl acetate, artemisia ketone and artemisia alcohol; Myrox: (*Z*)-myroxide, santolina triene and artemisyl acetate.

| | **Intercept** | **Chemotype** | **Plot type** | **Chemotype x Plot type** | **Day** | **Wind speed** | **Year** | **Flower head number**[a] | **Temperature** | **Humidity** | ***Block*** | ***Plot*** | ***Maternal origin*** | ***Plant*** |
|---|---|---|---|---|---|---|---|---|---|---|---|---|---|---|
| ***Apis mellifera* visits:** Zi-GLMM (Poisson distribution + log-link) | | | | | | | | | | | | | | |
| Type III Wald Chi-square tests | $\chi^2$ = 659.52<br>*df* = 1<br>***p* < 0.001** | $\chi^2$ = 1.72<br>df = 4<br>*p* = 0.797 | $\chi^2$ = 9.34<br>df = 1<br>***p* = 0.002** | $\chi^2$ = 25.87<br>df = 4<br>***p* < 0.001** | $\chi^2$ = 335.76<br>df = 1<br>***p* < 0.001** | $\chi^2$ = 0.85<br>df = 1<br>*p* = 0.356 | $\chi^2$ = 660.15<br>df = 1<br>***p* < 0.001** | $\chi^2$ = 100.761<br>df = 1<br>***p* < 0.001** | $\chi^2$ = 2.45<br>df = 1<br>***p*** = 0.118 | $\chi^2$ = 0.203<br>df = 1<br>*p* = 0.652 | -- | -- | -- | -- |
| Conditional model fixed effects levels or random effects variances | -- | Aacet – Intercept = - 0.026, n.s.<br>BThu – Intercept = 0.132, n.s.<br>Keto – Intercept = 0.036, n.s.<br>Myrox – Intercept = - 0.0125, n.s. | Homogeneous – Intercept = **- 0.315 **** | Aacet homogeneous – Intercept = **0.541 *****<br>BThu homogeneous – Intercept = 0.061 n.s.<br>Keto homogeneous – Intercept = **0.410**, ******<br>Myrox homogeneous – Intercept = **0.458 **** | -- | -- | -- | -- | -- | -- | *1.21 %* | *1.34 %* | *1.70 %* | *7.57 %* |
| Zero-inflation model | Estimate = - 1.43<br>***p* < 0.001** | NT | NT | NT | Estimate = 0.095<br>***p* < 0.001** | NT | NT | NT | NT | NT | NT | NT | NT | NT |
| **Phalacrid (primarily *Olibrus* spp.) visits:** Zi-GLMM (Poisson distribution + log-link) | | | | | | | | | | | | | | |
| Type III Wald Chi-square tests | $\chi^2$ = 0.12<br>df = 1<br>*p* = 0.734 | $\chi^2$ = 3.71<br>df = 4<br>*p* = 0.45 | $\chi^2$ = 0.013<br>df = 1<br>*p* = 0.911 | $\chi^2$ = 3.76<br>df = 4<br>*p* = 0.440 | $\chi^2$ = 15.618<br>df = 1<br>***p* < 0.001** | NT | $\chi^2$ = 0.1063<br>df = 1<br>p = 0.744 | $\chi^2$ = 5.06<br>df = 1<br>***p* = 0.025** | NT | $\chi^2$ = 29.85<br>df = 1<br>***p* < 0.001** | -- | -- | -- | -- |
| Conditional model fixed effects levels | -- | Aacet – Intercept = - 0.2485 n.s.<br>BThu – Intercept = | Homogeneous – Intercept = 0.061 n.s. | Aacet homogeneous – Intercept = - 0.213 n.s.<br>BThu homogeneous – | -- | -- | -- | -- | -- | -- | *2.21 %* | *1.56 %* | *0.43 %* | *11.47 %* |

| | | | | | | | | | | | | | | |
|---|---|---|---|---|---|---|---|---|---|---|---|---|---|---|
| or random effects variances | | - 0.00394 n.s.<br>Keto – Intercept = 0.101 n.s.<br>Myrox – Intercept = 0.0629 n.s. | | Intercept = - 0.289 n.s.<br>Keto homogeneous – Intercept = -0.349 n.s.<br>Myrox homogeneous – Intercept = - 0.046 n.s. | | | | | | | | | | |
| Zero-inflation model | Estimate = -2.049<br>***p* < 0.001** | NT | NT | NT | NT | NT | NT | Estimate = 0.122<br>***p* < 0.001** | NT | NT | NT | NT | NT | NT |

[a]: Flower head number used as covariate is freshly-opened flower heads for pollinators and mature (fresh or senescent) for florivore

### FIGURES

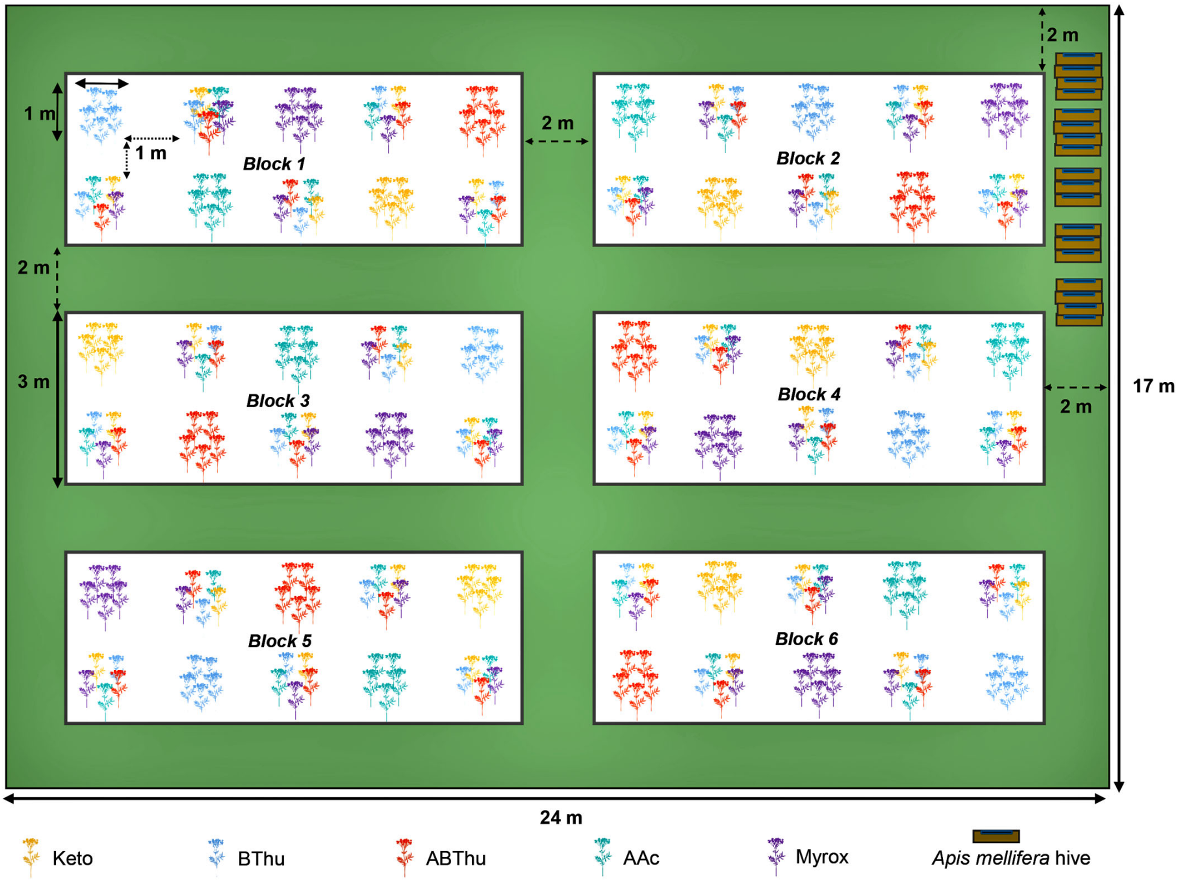


**Figure S1.** Design of field set-up showing different blocks and plots with the different *Tanacetum vulgare* chemotypes organised into homogeneous (single chemotype) and heterogeneous (mix of all five chemotypes) plots. (Keto: artemisia ketone chemotype, BThu: β-thujone chemotype, ABThu: α-/β-thujone chemotype, Aacet: artemisyl acetate/artemisia ketone/artemisia alcohol chemotype, Myrox: (*Z*)-myroxide/santolina triene/artemisyl acetate chemotype).

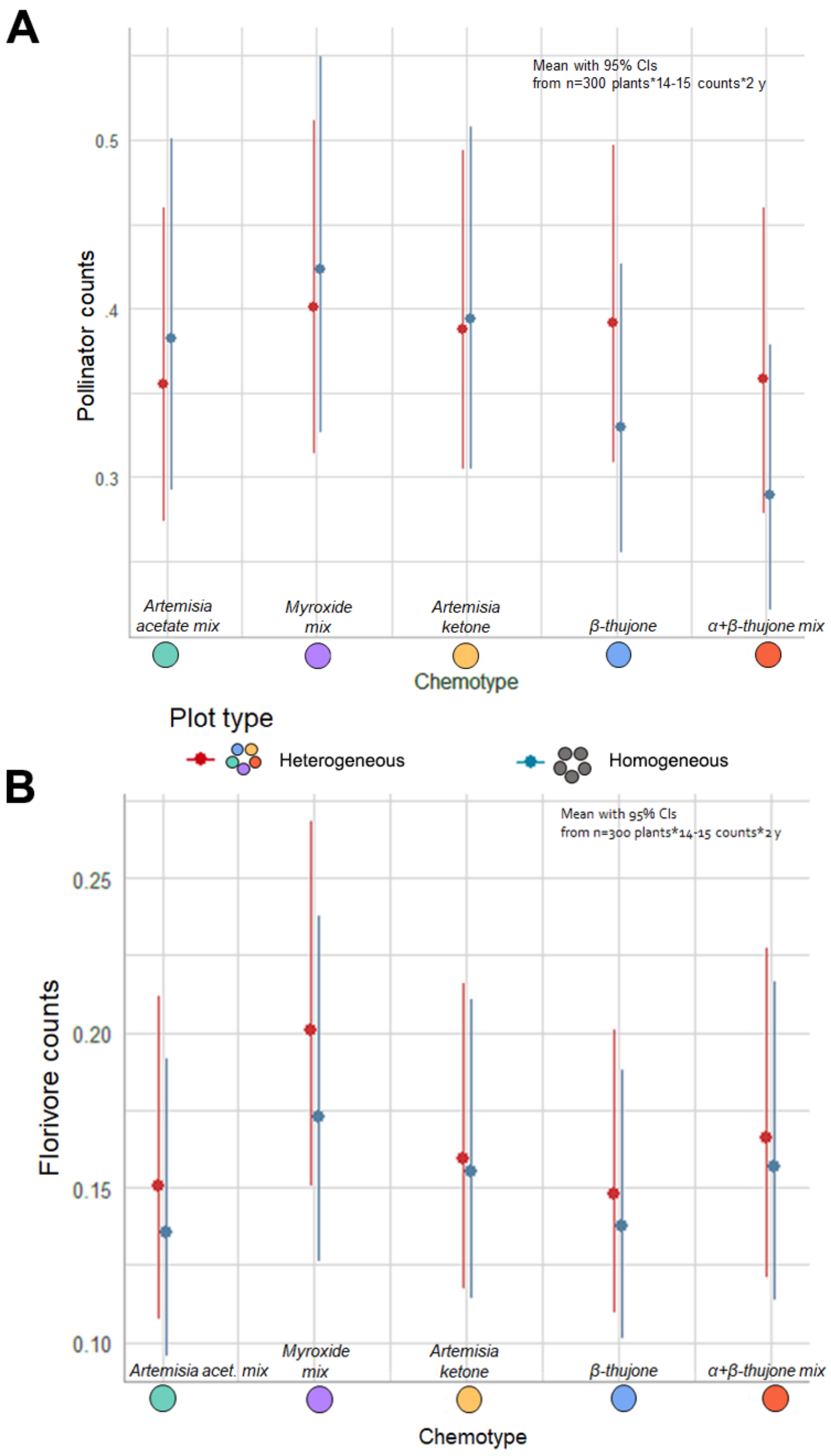


**Figure S2.** GLMM-estimated back-transformed means with 95% confidence intervals of (A) pollinator and (B) florivore counts in relation to chemotype (x-axis) and plot type.

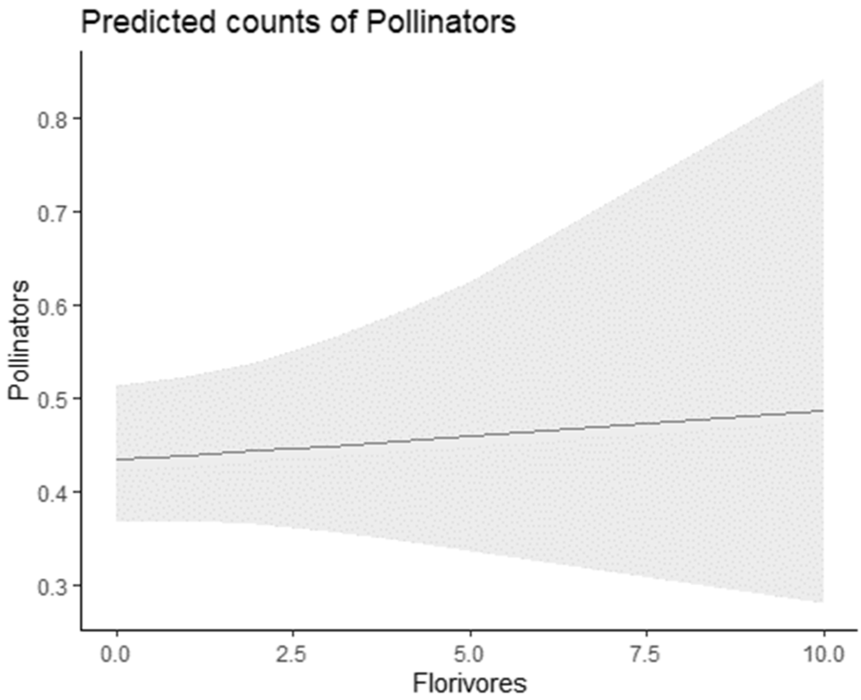


**Figure S3.** Relation between pollinators and florivores in 2022 (GLMM, $\chi^2 = 0.1744$, $p$ = 0.676).
